# Metabolic Reprogramming Induced by Mitochondrial Citrate Carrier Deletion Mitigates Antibiotics-Induced Acute Tubular Injury

**DOI:** 10.64898/2026.04.29.721583

**Authors:** Ming-Hung Hu, Kaihao Wang, Peir-In Liang, Elaine Y Dai, Adam J. Rauckhorst, Renny S. Lan, Hailemariam Abrha Assress, Eric Taylor, Dao-Fu Dai

## Abstract

**Introduction:** The mitochondrial citrate carrier (CiC), which mediates the transport of citrate across mitochondria, has been implicated in various diseases, but its role in kidney tubules is unclear. Here, we unraveled a novel role of CiC in tubular metabolism in the context of antibiotics-induced acute tubular injury (ATI).

**Methods:** ATI was induced by administration of vancomycin and gentamycin for 48 hours in mice (V+G-ATI). Tubular-specific CiC knockout (KO) was induced by adeno-associated virus (AAV) serotype 9 encoding Cre recombinase driven by KSP promoter (AAV9-Ksp-Cre) injection. Unbiased proteomic and metabolomic analyses were performed in CiC KO mouse kidneys. We performed *in vivo* ^13^C metabolic flux analysis to elucidate metabolic alterations in ATI and the effect of CiC KO.

**Results:** In this study, V+G-induced ferroptosis, oxidative damage, and extensive ATI in mice were alleviated by CiC KO. Metabolic reprogramming induced by CiC KO increased mitochondrial TCA cycle intermediates, including alpha ketoglutarate (AKG), and elevated levels of the endogenous antioxidant glutathione (GSH). Supplementation with AKG or GSH attenuated V+G-ATI in mice. Tracking of the ^13^C pyruvate / lactate revealed an increased flux of glucose oxidation pathway in V+G-ATI. Interestingly, tubular-specific CiC KO expands the effective TCA cycle pool reserve space, which may contribute to mitigation of ROS. The beneficial metabolic alteration in CiC KO requires AKG and glutamate, as simultaneous inhibition of mitochondrial transporters of AKG and glutamate attenuated the cytoprotective effects of CiC KO against antibiotic-induced oxidative damage.

**Conclusions:** This is the first study to demonstrate the role of mitochondrial CiC in kidney tubular epithelial cells, showing that it induces metabolic alterations that protect against antibiotic-induced ATI.

## Introduction

Acute kidney injury (AKI) is a common complication in critically ill patients, often leading to significant morbidity and mortality^1^. Medications are frequent causes of AKI, accounting for approximately 20% of all cases^2,3^. Among these, vancomycin is a widely used antibiotic that is particularly effective in treating methicillin-resistant Staphylococcus aureus bacteremia and infectious endocarditis^4^. The use of vancomycin and aminoglycosides (e.g. gentamicin) is often complicated by nephrotoxicity, which manifests as acute tubular injury (ATI)^5^. To date, there are no robust preventive strategies for antibiotic-induced ATI. This poses a significant challenge to balance the benefit of infection control versus the risk of AKI.

The kidney relies predominantly on mitochondrial bioenergetics to meet the substantial energy demands of solute reabsorption^6^. The citric acid (TCA) cycle, coupled with oxidative phosphorylation, constitutes the most efficient bioenergetic pathway for renal cells, particularly in proximal tubular epithelial cells. ATI is characterized by mitochondrial damage and metabolic alterations, with a shift in primary ATP production from the fatty acid to glucose utilization^7^. In recent years, emerging studies have highlighted the central role of mitochondrial dysregulation in the pathophysiology of ATI^8 9^.

The mitochondrial citrate carrier (CiC) encoded by the *SLC25A1* gene is a crucial mitochondrial membrane transporter that mediates the bidirectional transport of citrate across the mitochondrial membrane. Citrate plays a pivotal role in multiple metabolic pathways. It serves as a source of acetyl-Coenzyme A (Ac-CoA) for fatty acid and sterol biosynthesis, while also acts as a key regulatory molecule for glycolysis and gluconeogenesis.^10–12^ Previous studies showed that CiC is a critical regulator of TCA cycle that affects mitochondrial function and it has been implicated in various human diseases, including inflammation,^13^ neurodegeneration,^14^ and nonalcoholic fatty liver disease^15^. However, the role of CiC in kidney tubules is unclear.

In this study, we investigated the critical role of CiC in acute tubular injury (ATI) induced by a combination of vancomycin and gentamycin (designated as V+G-ATI). KEGG analysis of broad proteomic profiling data suggested that ferroptosis pathway is a critical mechanism of V+G-ATI. Ferroptosis is a form of reactive oxygen species (ROS)-driven cell death characterized by accumulation of iron leading to lipid peroxidation, such as through Fenton reaction.^6,16^ Free intracellular iron or iron-containing enzymes react with oxygen and polyunsaturated fatty acid (PUFA)-containing lipids, producing excessive membrane lipid peroxides, causing cell death.^17^ Tubular-specific CiC KO led to metabolic reprogramming that enhanced endogenous glutathione antioxidants. It restored GPX4 expression, leading to the inhibition of ferroptosis, and protected against V+G-ATI. The protective effect of CiC knock-down required the transfer of AKG and Glutamate across mitochondria.

Moreover, the administration of AKG or GSH, an endogenous antioxidant, conferred therapeutic benefit in ameliorating V+G-ATI, underscoring the importance of metabolic restoration in kidney protection.

## Materials and Methods

### Mouse experiments

CiC^flox/flox^ mice were generated and obtained as previously described.^12^ To generate CiC deletion (CiC ^−/-^) mice, nine 4-6-month-old CiC ^flox/flox^ mice of C57Bl6/J background (balanced gender) were treated with intraperitoneal injections of an adeno-associated virus (AAV) serotype 9 vectors encoding Cre recombinase driven by the KSP promoter (AAV9-KSP-Cre).^18^ This AAV delivery system facilitated the kidney tubules specific deletion of CiC. All mice were maintained under specific pathogen-free regulations at the Johns Hopkins University School of Medicine Animal Facility (Baltimore, Maryland, USA). All procedures were performed according to preapproved protocols and in accordance with recommendations for the proper use and care of laboratory animals.

### V+G ATI model

To create the antibiotics-induced ATI model, we injected 5-6 mice (both males and females) with Vancomycin (100 mg/kg/day) and Gentamicin (80 mg/kg/day) via intraperitoneal route for two consecutive days. Saline injection was performed for 3-5 mice as control groups. Blood and kidney samples were collected 48 hours after the first antibiotic dose. For the therapeutic metabolite intervention, α-ketoglutarate (AKG, 100 mg/kg) or glutathione (GSH, 100 mg/kg) was administered 24 hours and 4 hours prior to antibiotic treatment on both days. For the in vitro experiments, HK-2 cells were incubated with vancomycin (5 mM) and gentamicin (3 mM) in 4 mL of culture medium for the indicated durations.

### Cell culture

The human proximal tubular cell line, HK-2 (ATCC® CRL-2190™) was cultured in a 1:1 mixture of DMEM/F12 medium (Gibco, 11320-033) supplemented with: 10% fetal bovine serum (FBS), 15 mM HEPES, 50 nM hydrocortisone (Sigma, H-6909), 5 µg/mL insulin (Sigma, I-1882), 5 µg/mL transferrin (Sigma, T-0665), 50 ng/mL sodium selenite (Sigma, S-9133), 1 ng/mL 3,3′5-triiodo-L-thyronine (T3, Sigma, T-5516), and 10 ng/mL epidermal growth factor (BD Biosciences, 354001). Cells were maintained in an atmosphere of 95% air and 5% CO₂ at 37°C

Additional details for all methods are provided in Supplementary Methods.

## Results

### CiC KO exerts a protective effect against vancomycin and gentamycin-induced acute kidney injury in mice

Figure 1A summarizes the mouse experimental design. The mitochondrial citrate carrier (CiC ^flox/flox^) mice at 4–6-month-old received Adeno-associated virus (AAV) serotype 9 vectors encoding Cre recombinase driven by the KSP promoter (AAV9-KSP-Cre) by intraperitoneal injection to induce CiC deletion in kidney tubules. Approximately 2 weeks after induction of kidney tubules specific CiC KO, ATI was induced by daily intraperitoneal injections of V+G for two consecutive days (Fig. 1A). Gentamycin was added to potentiate the nephrotoxic effect of vancomycin. Kidney tissues were harvested 48 hours later. V+G injection in wild type (WT) mice led to elevated serum BUN levels, and extensive pathological ATI, characterized by significant vacuolation, attenuation of tubular epithelial cells, tubular dilation, and cell death, compared with saline controls (Fig 1B, C). Quantitative pathological analysis showed that approximately 50% of tubules were injured in WT mice 48 hours after V+G. Notably, tubular specific CiC KO mice exhibited substantial protection, with significantly lower serum BUN and less tubular injury compared to WT V+G (Fig 1D). These findings indicate that inhibition of CiC ameliorated V+G-ATI.

**Figure 1.**
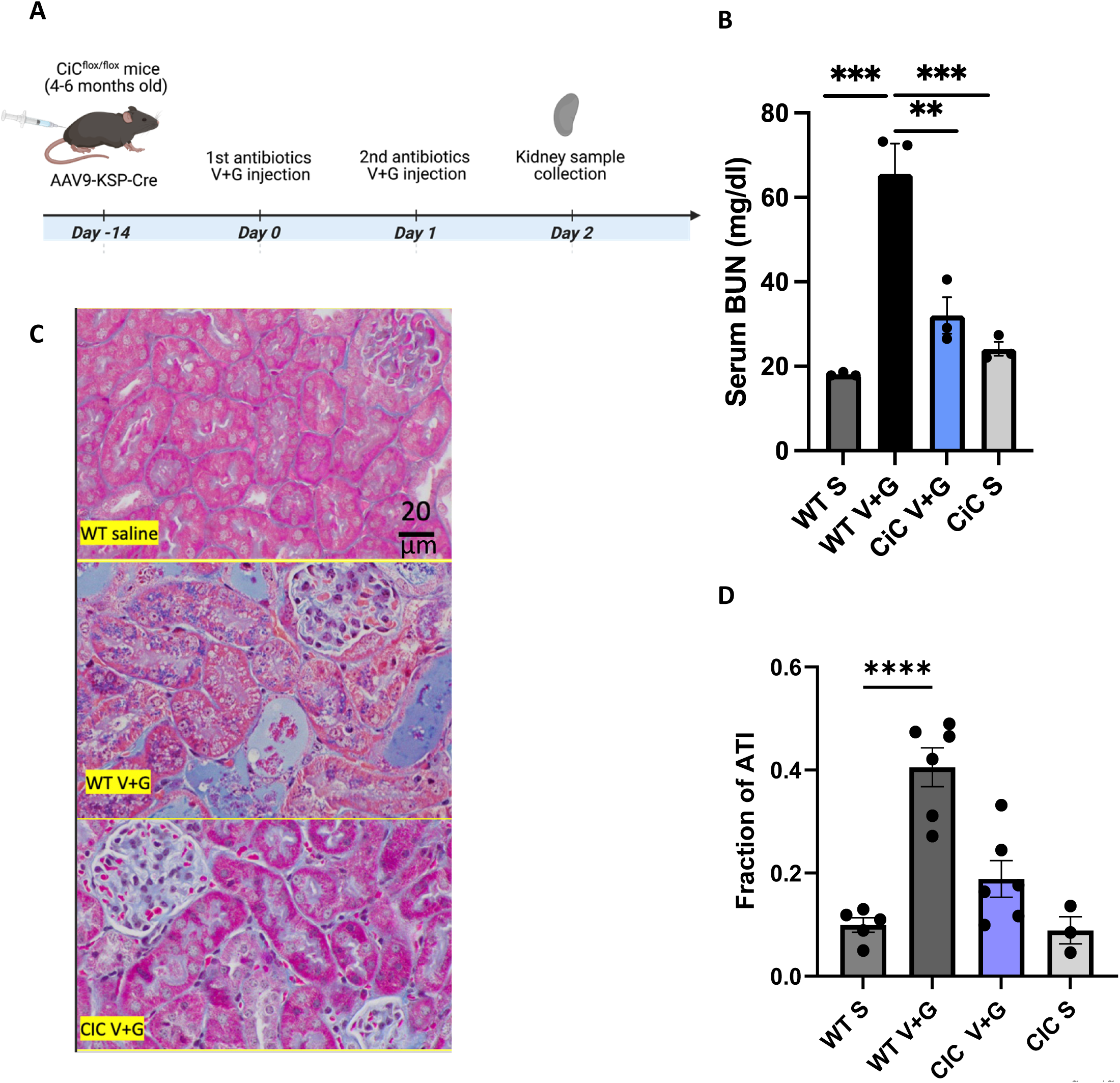
CiC knockout mitigates antibiotics-induced acute kidney injury in mice. **(A)** Experimental design. CiC^flox/flox^ mice were treated with AAV9-KSP-Cre via intraperitoneal injection to induce CiC deletion in kidney tubules, followed by injection of vancomycin and gentamicin (V+G) for 2 days. (**B)** Serum BUN levels 48 hours after V+G. **(C)** Representative Trichrome stain sections showed acute tubular injury after V+G. **(D)** Fraction of acute tubular injury. 2-way ANOVA. Sidak post-hoc tests for 2-group comparisons, **, *p*< 0.01; ***, *p* < 0.001; ****, p<0.0001. scale bar: 20 µm. S: Saline; CiC: Citrate Carrier knock-out (tubular specific)

### Unbiased proteomic profiling and KEGG pathway analysis highlights the ferroptosis pathway in vancomycin/gentamicin induced ATI

To investigate the plausible molecular mechanisms underlying acute kidney tubular injury in the V+G-ATI mouse model, we performed an unbiased quantitative proteomic profiling by labeled-free Data Independent Analysis (DIA) (supplement method) (Supplement Data 1) (Fig S1). Compared with saline controls, the WT mice receiving V+G show 320 proteins significantly altered (FDR-adjusted p-values <0.05). Using the cut-off of at least 2-fold change (log2 fold change of +/− 1), the volcano plots summarize the differential protein expression analysis of the proteomic profiling data. The plot shows that 229 proteins were significantly upregulated, and 91 proteins were significantly downregulated in V+G-ATI in WT mice, suggesting a substantial impact of antibiotics-induced ATI on the kidney proteome (Fig 2A, B) (Fig S2A, B).

**Figure 2.**
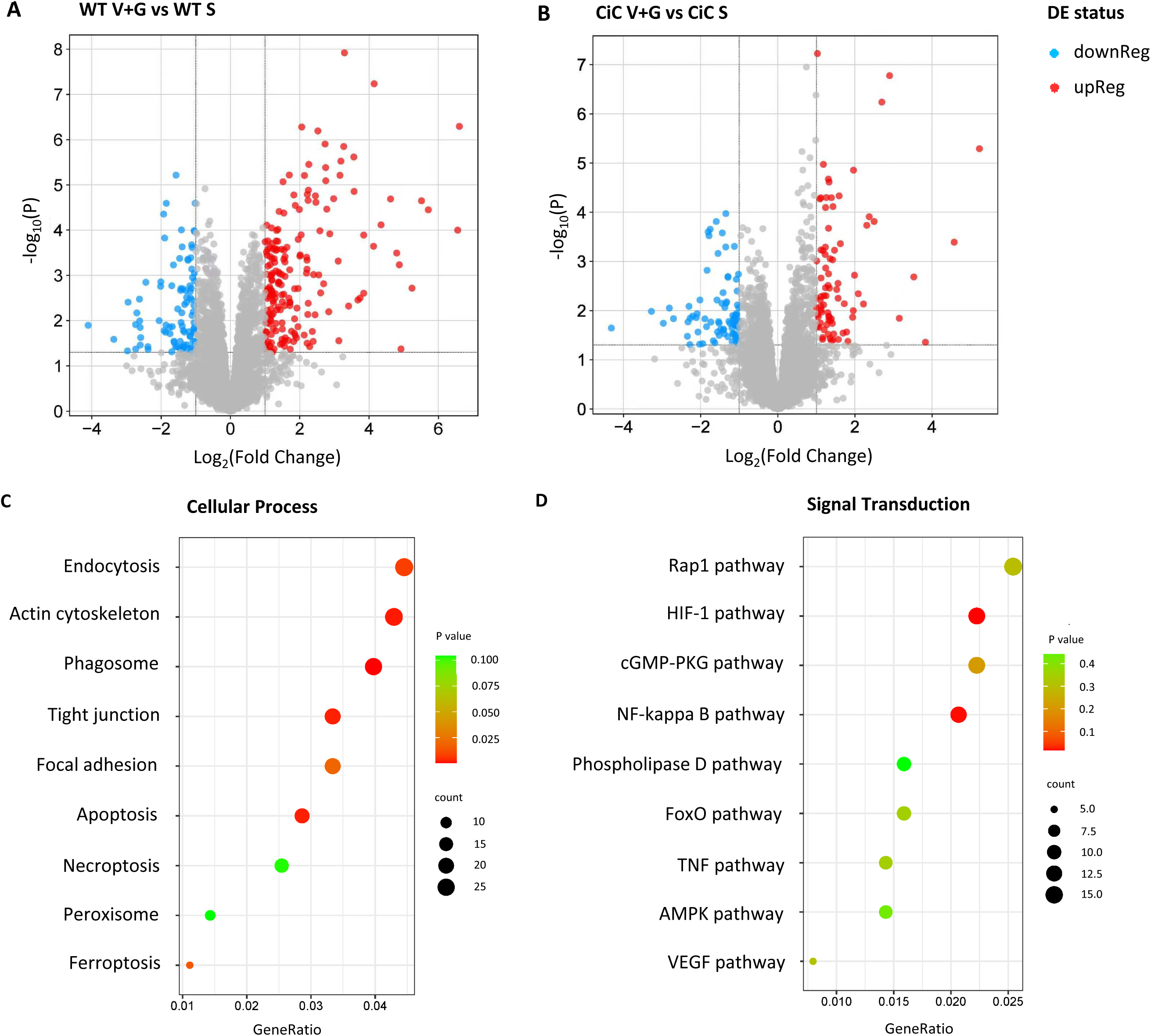
KEGG pathway analysis of proteomics data identifies alterations in the ferroptosis pathway during antibiotics-induced acute tubular injury. Unbiased proteomics analysis was performed using label-free Data Independent Acquisition. **(A)** Volcano plot showing 320 proteins significantly changed between the V+G-ATI and saline-treated control wild type (WT) mouse kidneys (fold change > 2, p < 0.05). Limma with empirical Bayes moderation, followed by FDR adjustment (Benjamini–Hochberg), with significance declared at FDR-adjusted p < 0.05 and absolute fold change > 2. (B) Volcano plot showing 173 proteins significantly changed between the V+G-ATI and saline-treated control CiC-KO mice (fold change > 2, p < 0.05). KEGG pathway enrichment analysis was performed for significant biological pathways in the CiC-KO versus V+G-ATI WT. (C) The enriched pathways of cellular process include endocytosis, cytoskeletal regulation, and ferroptosis, suggesting metabolic reprogramming associated with renal tubular-specific CiC KO. **(D)** KEGG pathways highlighting major signal transduction processes. DE: differentially expressed, either upregulation, downregulation or non-significant. Gene ratio is defined as the number of genes from the input list annotated to a given KEGG pathway divided by the total number of genes in the input list used for enrichment analysis.

The perturbative effect of V+G was markedly blunted in CiC KO mice, with only 90 proteins being upregulated and 83 downregulated. To gain insight into the underlying mechanisms, we performed the KEGG (Kyoto Encyclopedia of Genes and Genomes) pathway analysis comparing WT V+G vs CiC V+G. Significantly overrepresented cellular processes include phagosome, apoptosis, and ferroptosis pathways (Fig. 2C). The HIF-1 and NF-_κ_B were the overrepresented and significantly altered signal transduction pathways (Fig.2D). Furthermore, amino acid metabolism overrepresentation highlighted branched-chain amino acid degradation, cysteine, methionine, alanine, aspartate and glutamate metabolisms (Fig S3). Together, these findings suggest the critical pathways involved in V+G-ATI include cell death (apoptosis, ferroptosis), inflammatory and hypoxia pathways, and amino acid metabolism.

### CiC KO mice modulate GPX4, and other critical proteins involved in ferroptosis

Ferroptosis is a form of cell death characterized by iron-dependent lipid peroxidation Our proteomics data reveal that several crucial proteins in the ferroptosis pathway were significantly altered in V+G-ATI. Glutathione Peroxidase-4 (GPX4), a key ferroptosis regulator, was lower in the V+G-ATI in WT but was relatively preserved in tubular-specific CiC KO mice, as shown by quantitative proteomic analysis (Fig. 3A). The other well-known markers of ferroptosis are elevated levels of Acyl-CoA synthetase long-chain family member 4 (ACSL4) and transferrin receptor protein (TFRC). Proteomic analyses showed that both ACSL4 and TFRC were significantly elevated in V+G-ATI in WT, but more mildly and not significant in V+G-ATI in CiC KO (Fig. 3B, C). Western blot analysis showed significant changes in GPX4, ACSL4, and TFRC in V+G-ATI in CiC KO versus WT mice, consistent with the proteomic data (Fig. 3D-G). These findings suggest that tubular-specific CiC KO restored GPX4 and attenuated ferroptosis induced by V+G-ATI.

**Figure 3.**
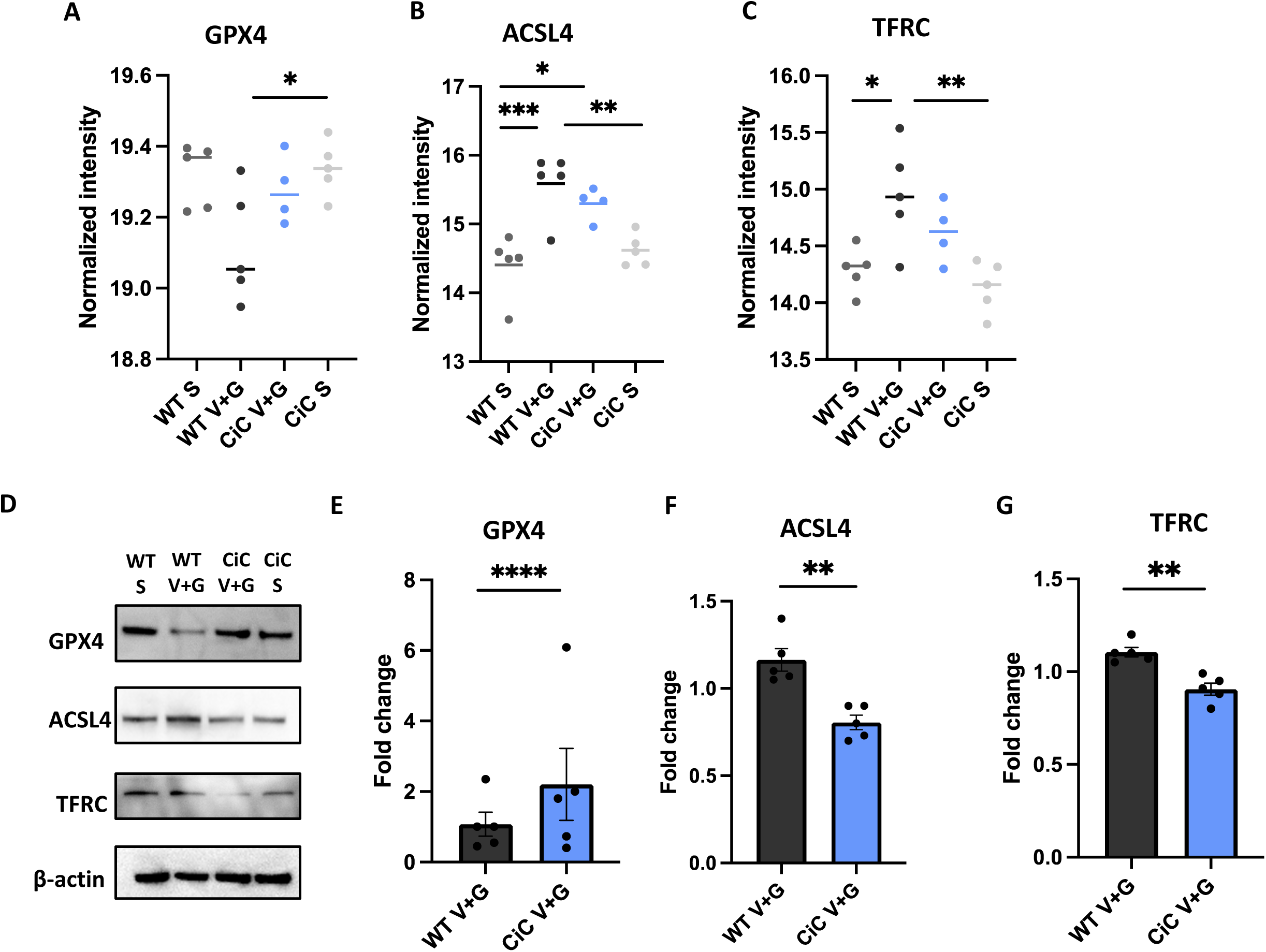
CiC knockout modulates GPX4 and other key regulators of the ferroptosis pathway in mouse kidneys. Quantitative proteomic analysis of **(A)** GPX4 (Glutathione Peroxidase-4), **(B)** ACSL4 (Acyl-CoA synthetase, long-chain family member 4), and **C)** TFRC (Transferrin receptor) levels determined by Data Independent Analysis in WT and tubular-specific CiC KO mouse kidney lysates. **(D)** Western blot of GPX4, ACSL4, and TFRC in WT and CiC KO kidneys. **(E-G)** Quantification of GPX4, ACSL4 and TFRC protein levels by Western blots. ANOVA post-hoc tests for A-C; two sample t-tests for E-G, *, *p* < 0.05; **, *p*< 0.01; ***, p<0.001, ****, *p* < 0.0001. S: Saline; CiC: Citrate Carrier KO (tubular specific)

### CiC knockdown (KD) attenuates oxidative stress and ameliorates ferroptosis

To evaluate the role of ferroptosis in V+G-ATI, we performed cell experiments using HK-2 human proximal tubular cell lines. We labeled cells with MitoSOX™ and DCFDA to assess mitochondrial ROS and total cellular H_2_O_2_ using flow cytometry. In HK-2 cells, the V+G increased MitoSOX (mitochondrial ROS) and DCFDA fluorescence (total H_2_O_2_, Fig. 4A-B). Prolonged oxidative stress may lead to lipid peroxidation, a crucial step of ferroptosis. We analyzed C11-BODIPY581/591™ fluorescence and showed a significantly higher lipid peroxidation level in V+G-treated HK-2 cells compared to saline controls (Fig. 4C). Knock down of CiC by siRNA in HK-2 cells resulted in significantly lower lipid peroxidation (Fig 4D), and total ROS in V+G (Fig 4E). In a complementary experiment, the selective CiC inhibitor CTPI-2 recapitulated these effects and significantly decreased mitochondrial and total ROS (Fig 4FG). Moreover, GPX4 levels were better preserved in CiC KD HK-2 cells (Fig S4).

**Figure 4.**
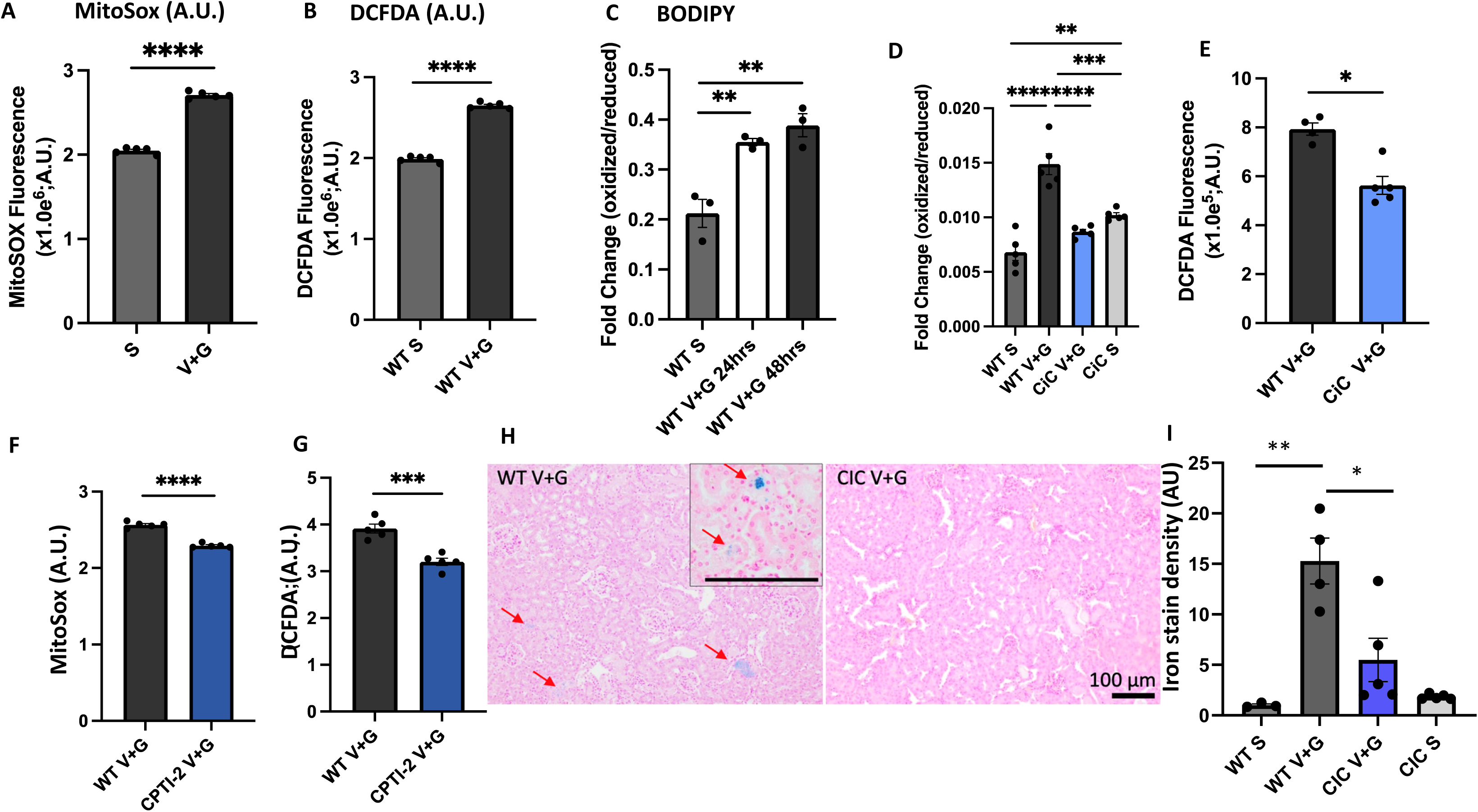
Amelioration of V+G-ATI by CIC KO via ferroptosis suppression. MitoS^TM^ Red and DCFDA fluorescence were quantified by flow cytometry to assess mitochondrial and total cellular oxidative stress, following one hour of V+G treatment in HK2 cells. **(A)** MitoSOX and **(B)** DCFDA fluorescence increased in response to V+G. **(C)** Lipid peroxidation analysis using BODIPY^TM^ revealed significantly elevated lipid peroxidation levels in V+G-treated HK-2 cells compared to controls. The increased oxidative stress in response to V+G treatment, **(D)** MitoSox and **(E)** DCFDA, were significantly attenuated in CIC-knock-down HK-2 cells. Selective CIC inhibitor, CPTI-2 (50 µM for 3 hours) significantly attenuated **(F)** MitoSox and **(G)** DCFDA in HK-2 cells challenged with V+G. **(E)** Representative images of iron stain showed accumulation of iron (blue, red arrows) in injured tubules. **(F)** Relative density of iron staining. A.U. Arbitrary unit of fluorescence or color intensity. ANOVA post-hoc tests or two-sample t-tests (*, *p* < 0.05; **, *p*< 0.01; ***, *p*< 0.001; ****, *p*< 0.0001). S: Saline; CIC: citrate carrier siRNA or KO

Given data in HK-2 cells consistent with CiC protecting from ferroptosis, we extended our evaluation of ferroptosis in mouse kidneys. Iron staining of kidney sections showed scattered tubular epithelial cells with accumulation of iron, another hallmark of ferroptosis, particularly within injured tubules in V+G-ATI. As shown in Fig 4H-I, there was substantial accumulation of iron in V+G-ATI in control mouse kidneys (CiC ^fl/fl^ mice treated with AAV9 vector only, no deletion). In contrast, CiC KO exhibited a significant amelioration in iron accumulation within kidney tubules (60% less by relative density). Collectively, these findings suggest that V+G induced ferroptosis, and that CiC KD ameliorated ferroptosis and decreased iron-dependent lipid peroxidation.

### Metabolic reprogramming by CiC KO induced a distinct metabolic profile

Metabolomics was performed to elucidate the metabolic reprogramming by tubule-specific CiC KO in the context of V+G ATI (Fig S5) (supplement method). Unbiased methods identified a total of 632 metabolites. Among these, 103 metabolites (50 upregulated and 53 down-regulated) significantly changed in ATI (V+G-ATI vs saline control). This effect was blunted in CiC KO mice, decreasing to 65 metabolites (29 upregulated and 36 down-regulated) significantly modified by CiC (CiC vs WT, both injured by V+G, Fig 5A) (Supplement Data 2, 3). The metabolomics analysis was visualized using a cnetplot (Fig. 5B) (Table 1). Among these, 91 and 71 metabolites matched well-annotated HMDB compounds in MetaboAnalyst 6.0. The enrichment and pathway analyses were performed against both KEGG (Fig 5C) and SMPDB libraries using Fisher’s Exact Test (Supplement Data 4, 5, Fig S6, 7). Metabolite enrichment is expressed as the fractional abundance of the input list relative to total metabolite pool. This network-based representation effectively demonstrates how key metabolites are shared across multiple biological processes, providing insights into functional linkages and potential regulatory hubs. The heat map from the CiC KO group showed a distinctive pattern between WT and CiC KO, as represented by red and blue colors (Fig. 5D). Furthermore, a bubble plot revealed the top 50 protein and metabolite change (p< 0.05), highlighting relationships between significantly enriched metabolic pathways and their associated metabolites (Fig. 5E). Our results demonstrated a distinct metabolic profile induced by the renal tubule-specific CiC KO condition.

**Figure 5.**
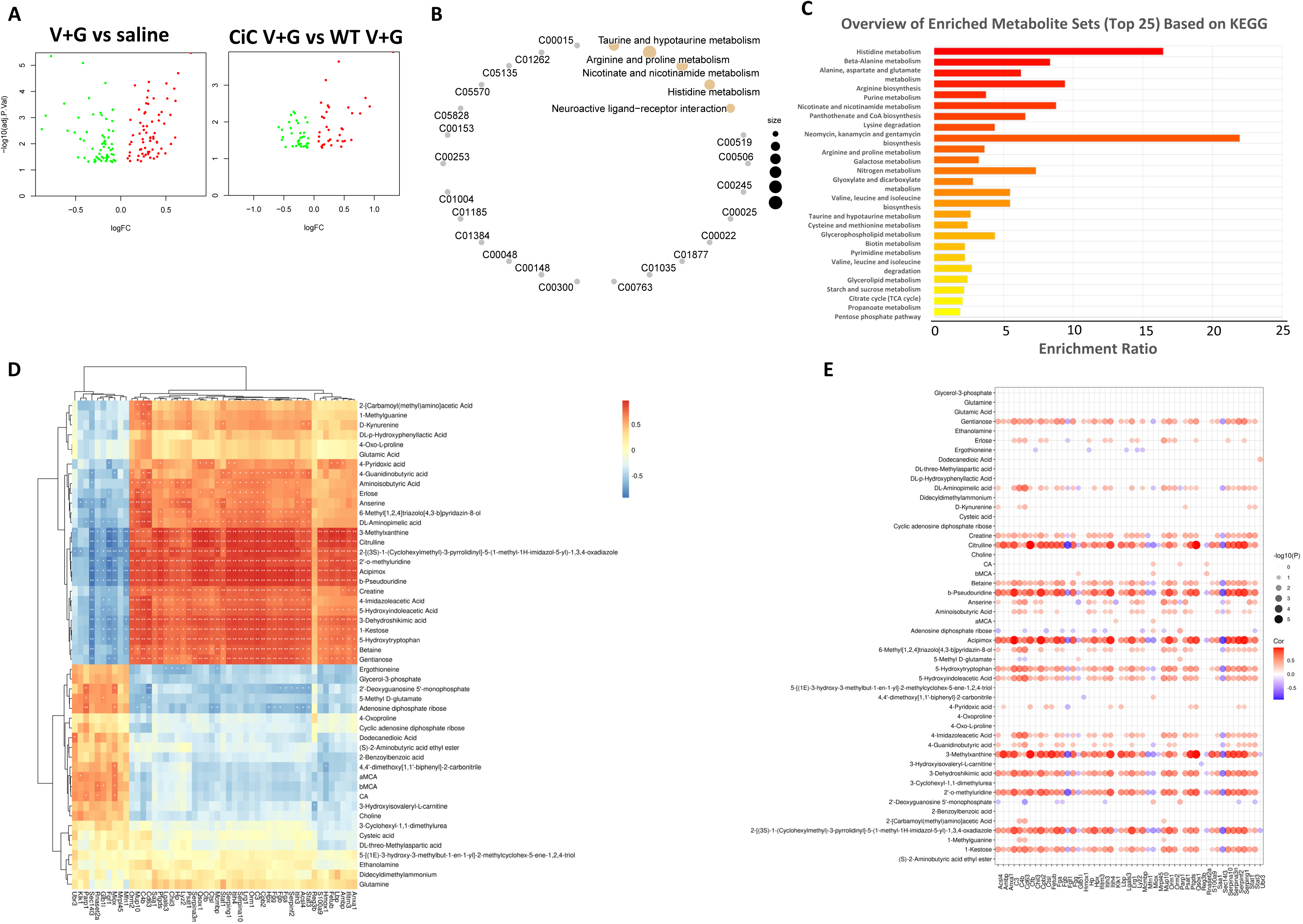
Metabolomics analysis identifies unique metabolite changes in tubular-specific CiC-KO and V+G ATI. Untargeted metabolomics analysis of snap-frozen mouse kidneys. **(A)** Volcano plot analysis revealed that 65 metabolites changed significantly between V+G-treated CiC-KO and wild-type mice. Similarly, 103 metabolites were significantly altered between V+G-treated and saline-treated control mice (fold change > 2, p < 0.05). Limma with empirical Bayes moderation, followed by FDR adjustment (Benjamini–Hochberg), with significance declared at FDR-adjusted p < 0.05 and absolute fold change > 2. **(B)** The cnetplot visualizes the relationships between significantly enriched metabolic pathways and their associated metabolites identified in CiC KO vs. control mice. **(C)** KEGG-based pathway enrichment analysis displays significantly enriched KEGG pathways derived from metabolites differentially expressed between CiC KO and control groups. The y-axis represents pathway names, and the x-axis indicates enrichment ratio. **(D)** Heatmap illustrates the relative abundance of significantly altered metabolites across CiC KO and control mice. Color intensity reflects normalized expression values, with red indicating upregulation and blue indicating downregulation. Hierarchical clustering was applied to both rows and columns using Euclidean distance and average linkage to reveal sample groupings and metabolic trends. **(E)** The bubble plot displays the top enriched metabolic pathways based on KEGG analysis of differentially expressed metabolites. Bubble size corresponds to the number of matched metabolites per pathway, and the color gradient reflects the statistical significance (adjusted p-value), with darker shades indicating stronger significance.

### Mitochondrial transporters for AKG and glutamate are required for the cytoprotective effect of CiC inhibition against antibiotics-induced oxidative stress

Our data showed that inhibition of mitochondrial citrate carrier leads to metabolic reprogramming that confers resistance to V+G-induced ATI. Because CiC function is expected to affect TCA metabolites, we summarize the metabolomics data and highlight the ratio of mouse kidney metabolites in V+G-ATI (WT V+G / WT saline, green) and the effect of renal tubule-specific CiC KO (CiC KO V+G / WT V+G, blue) (Figure 6A). We focus on the TCA-GSH pathway and summarize the fold-change of metabolites in response to V+G-ATI and the alterations by CiC. In parallel with the reno-protective effect of CiC KO against V+G-ATI, the CiC-KO V+G had significantly higher reduced GSH compared with WT V+G. Consistent with this, there were ∼25% increases in AKG and Cysteine (the precursors of GSH). In WT mice, a significant increase of ∼27% in oxidized GSH (GSSG) in response to V+G-ATI is also consistent with the findings of increased oxidative stress, iron-accumulation, and the subsequent iron-induced lipid peroxidation, together with several other markers of ferroptosis. Overall, our data suggest the alteration of the TCA–GSH pathway induced by V+G-ATI and how they were modified by CiC KO.

**Figure 6.**
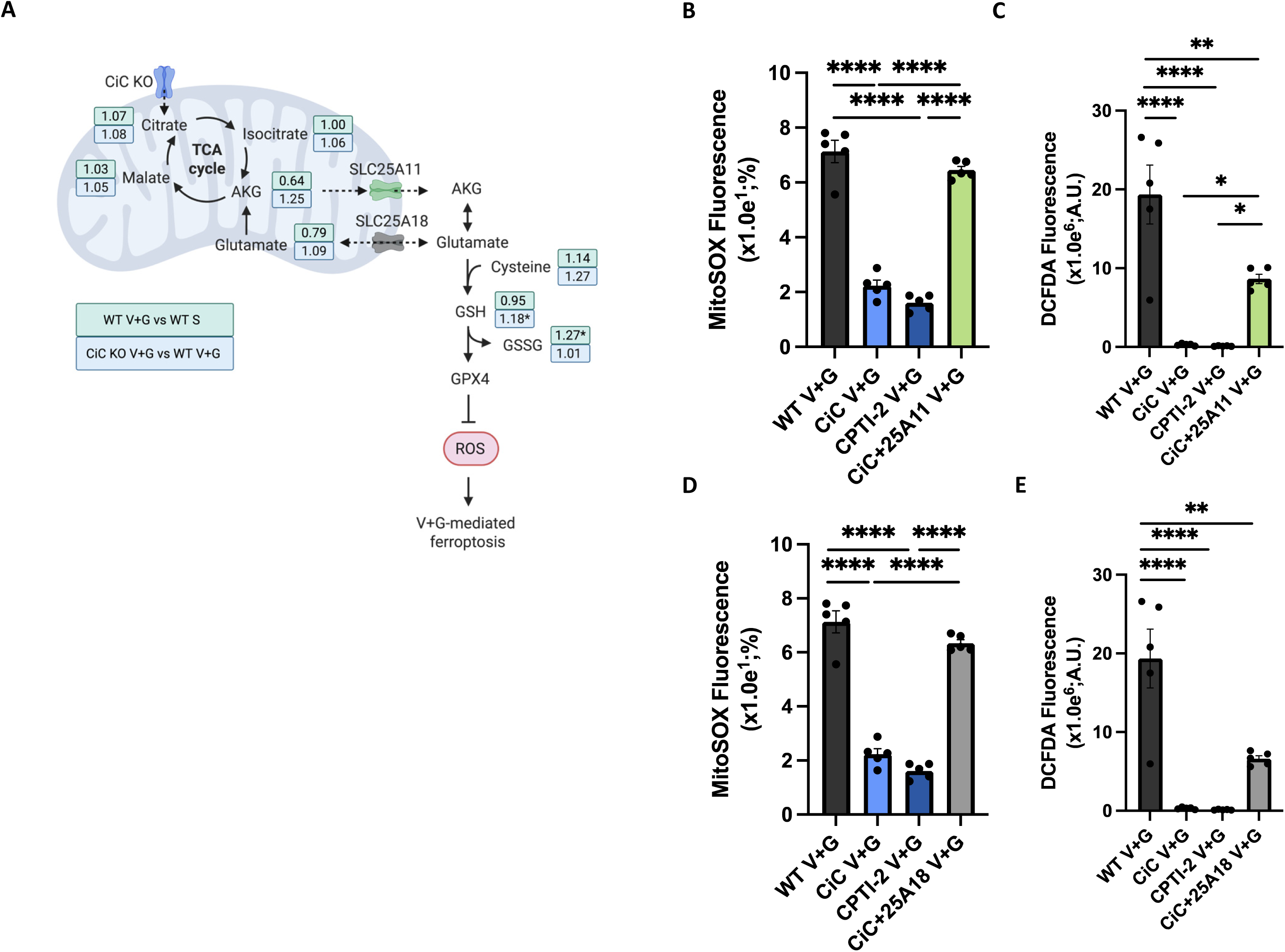
AKG and Glutamate trafficking play critical roles in mediating protection against V+G-induced oxidative stress. **(A)** Summary of the ratios of representative metabolites. Green boxes: WT V+G vs WT saline (the effect of ATI); Blue boxes: CiC V+G vs WT V+G (the effect of CiC KO). Several TCA cycle intermediates, along with GSH, were elevated in CiC KO. **(B-E)** The siRNA knock-down of SLC25A11 (AKG transporter) or SLC25A18 (Glutamate transporter) abolished or attenuated the protective effect of CiC inhibition. (B) MitoSox and (C) DCFDA fluorescence (arbitrary units) in HK-2 cells challenged by V+G, and the effect of CiC knock-down/inhibition, with or without concomitant suppression of SLC25A11. (D) MitoSox and (E) DCFDA fluorescence (arbitrary units) in HK-2 cells challenged by V+G, and the effect of CiC knock-down/inhibition, with or without concomitant suppression of SLC25A18. ANOVA post-hoc tests, (*, p<0.05, **, *p*< 0.01; ****, *p*< 0.0001).

Since GSH is primarily synthesized in cytosol, we hypothesize that mitochondrial transport of metabolites is essential for linking AKG metabolism to GSH biosynthesis. To examine the role of the AKG transporter in this context, we suppressed *SLC25A11*, the mitochondrial AKG transporter, using siRNA. Compared to sole CiC inhibition by siRNA knock-down or CTPI-2–treated HK-2 cells, cells with concomitant knock-down of *SLC25A11* exhibited significantly higher levels of mitochondrial and total ROS. Thus, suppression of the AKG transporter abolished the cytoprotective effect of CiC inhibition (Fig. 6B, C). Likewise, knock-down of the mitochondrial glutamate transporter, *SLC25A18* in CiC -KD HK-2 cells also attenuated the cytoprotective effect of CiC-KD against V+G-induced oxidative stress (Fig. 6D, E). These findings suggest that transfer of AKG as well as glutamate across mitochondrial membranes is essential for the CIC-mediated protective effect. Taken together, AKG and GSH are key metabolites involved in CiC KO-mediated metabolic reprogramming that alleviates antibiotics-induced oxidative stress, which is coordinated with AKG and glutamate transport.

### AKG or GSH supplementation ameliorate V+G induced ATI

To elucidate what aspect of metabolic reprogramming induced by CiC KO is cytoprotective, we focused on two metabolites that were increased in CiC KO: AKG, a TCA intermediate that has pleiotropic benefits^19^, and the endogenous antioxidant GSH, which is also enhanced by AKG^20^. HK-2 cells were pre-treated with AKG (8mM) or GSH (8mM), 24 hours prior to stressed by V+G. Consistent with antioxidant effects, either AKG or GSH attenuated mitochondrial and total cellular ROS induced by V+G (Fig 7A, B). To confirm this *in vivo*, we injected AKG (100mg/kg) or GSH (100mg/kg), initiated one day prior to V+G challenge in WT mice (Fig. 7C). The severity of acute tubular injury was protected by either GSH or AKG, as shown by significantly lower serum BUN (Fig 7D), and the fraction of injured tubules by pathology (Fig. 7E, F). To summarize, this *in vivo* experiment confirmed that key metabolites derived from the TCA cycle exert a protective effect against V+G-ATI, further highlighting the crucial role of metabolic interplay in mitigating drug-induced acute tubular injury.

**Figure 7.**
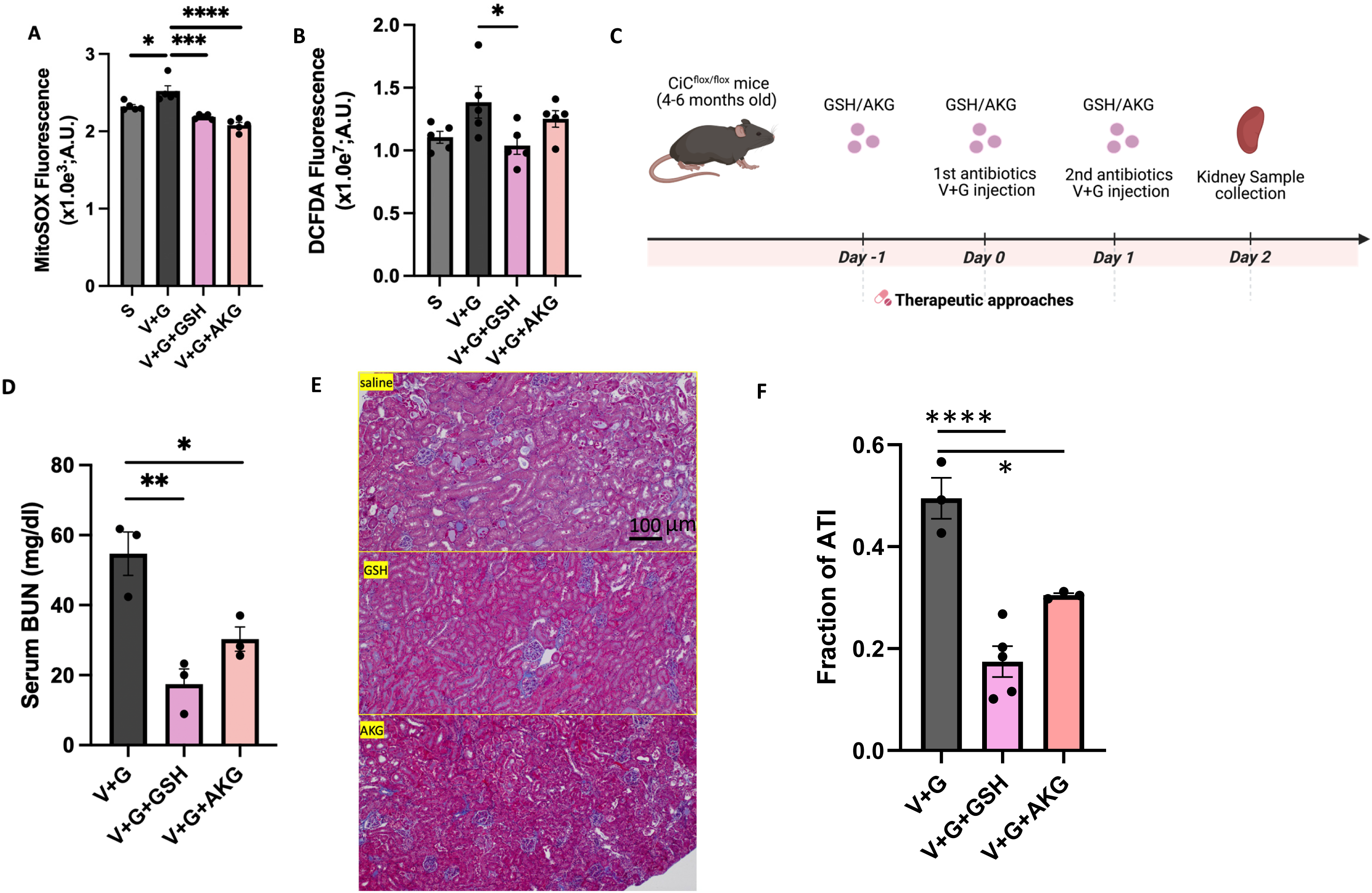
Supplementation with AKG or GSH attenuated V+G induced oxidative stress and acute tubular injury. **(A, B)** MitoSox and DCFDA fluorescence were significantly decreased in AKG or GSH–treated HK-2 cells challenged with V+G. **(C)** Experimental design showed wild type mice (CiC flox without Cre) receiving V+G stress, with or without prior AKG or GSH injection, started at 24h prior to V+G. **(D)** Serum BUN levels (48 hours post V+G). **(E, F)** Representative trichrome images and quantitative analysis of fraction of acute tubular injury. 2-way ANOVA post-hoc tests, (*, *p* < 0.05; **, *p*< 0.01; ***, *p*< 0.001; ****, *p*< 0.0001). S; Saline; AKG: α-Ketoglutarate; GSH: Glutathione.

### Increased glucose oxidation pathway in V+G-induced acute kidney injury

To explore possible switch in substrate utilization during acute kidney injury induced by V+G, we conducted ^13^C metabolic flux analysis to track the metabolites feeding into TCA cycle. We injected mice with uniformly labeled [U-¹³C]-lactate and [U-¹³C]-pyruvate stable-isotope tracers. ^13^C-enrichments into downstream metabolites were assessed 30 minutes after injection of labeled substrates in WT and CiC KO mice with V+G-ATI (Fig. 8A). TCA cycle M+2 ^13^C-enrichments denoting oxidizing, forward TCA cycle flux through acetyl-CoA, and were analyzed to isolate the mitochondrial oxidative fate of glucose (Fig. 8B). To evaluate the complex interplay between V+G-ATI and the effect of CiC KO, we performed multivariate regression analysis to evaluate the association between V+G-ATI or genotype (CiC KO) and the percentage of ^13^C-labeled metabolites. The multivariate regression analysis revealed that V+G-ATI significantly predicted many ^13^C metabolite M+2 enrichments versus the uninjured saline treated kidneys. In response to V+G-ATI, nearly all TCA intermediates, including citrate, isocitrate, AKG, succinate, fumarate, and the overall TCA pool, showed a robust increase in ^13^C-enrichment, as shown by positive beta values (Fig. 8C).

**Figure 8.**
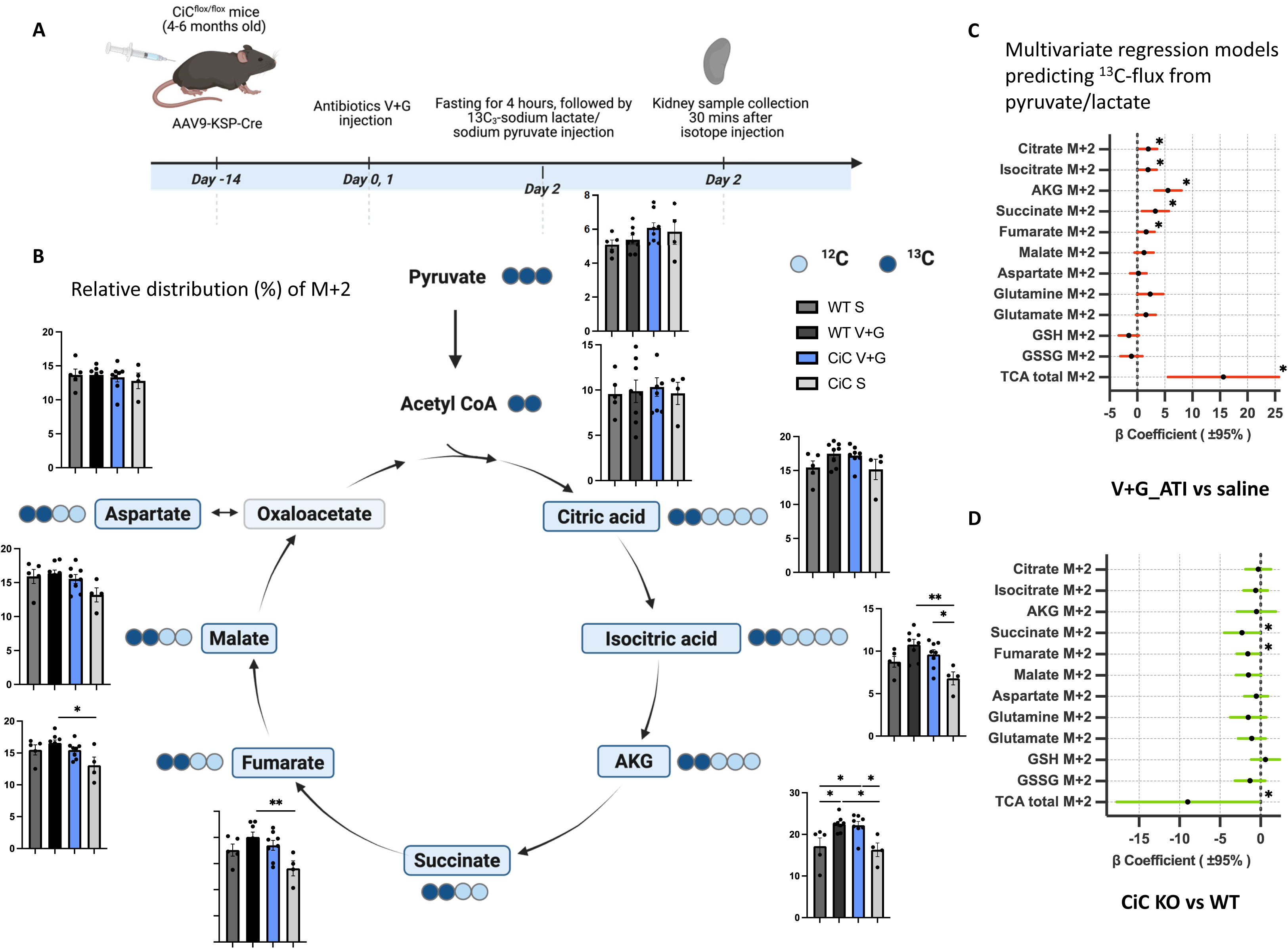
Metabolic Flux Analysis. [U-¹³C]-lactate and [U-¹³C]-pyruvate isotope tracing experiments were performed to investigate the CiC KO-mediated reno-protective pathway. (A) Schematic diagram of isotope tracing experiment design. Metabolite incorporation was assessed 30 minutes after isotope injection. (B) Two ^13^C (dark blue) from Acetyl CoA incorporated into TCA cycle. The relative distribution of labeled TCA intermediate metabolites (M+2) was presented in bar graphs. (C) Forest plot showing the results of the multivariable regression model, predicting M+2 (flux) from pyruvate/lactate. Each square represents the estimated coefficient predicting the metabolite, and the horizontal lines indicate the 95% confidence intervals. All data were presented as the mean ± SEM, (*, p < 0.05; **, p< 0.01).

In contrast, CiC KO had negative beta values (significant only for succinate, fumarate, and total TCA) (Fig. 8D). A mild decrease in M+2 fluxes in saline-treated CiC KO mice suggests increased TCA pool size reserve space, which permits greater glucose oxidation during V+G while limiting ROS induction that could contribute to ATI. Together with our other results, these isotope tracing results suggest that under antibiotic-induced stress, renal tubular cells upregulate glucose oxidation with potential ROS costs^8^, highlighting a metabolic vulnerability in the adaptive response to acute injury. CiC KO may mitigate this vulnerability without impairing adaptation, thereby protecting against oxidative damage, ferroptosis, and tubular injury.

## Discussion

To the best of our knowledge, the current study is the first to genetically test the role of mitochondrial CiC in kidney tubular epithelial cells *in vivo*. CiC KO did not affect kidney function or histology (at baseline) but it induced substantial metabolic alterations, and it also affected some protein (34 upregulated; 42 downregulated). Our data showed that, in response to vancomycin/gentamicinlZinduced ATI, CiC KO increased AKG levels and upregulated the GSH oxidative defense system, thereby altering metabolic flux in parallel with attenuated oxidative damage and ferroptosis. We show that injection of these metabolite supplements recapitulated the nephroprotective effect against V+G-ATI.

CiC facilitates the exchange of mitochondrial citrate for cytosolic malate. Efflux of citrate from mitochondria to cytosol have been shown to play important roles in inflammation, cancer cell reprogramming, histone acetylation, and insulin secretion ^21–23^. Previous study using a third-generation CiC inhibitor, CTPI-2,^24^ showed that it reversed hepatic steatosis, in addition to significant reduction of obesity, serum cholesterol, triglyceride, as well as glucose levels ^12,15^. Additionally, CiC downregulation mitigates inflammatory activation.^25^ CiC inhibition has also been shown to suppress the growth of various cancer types^24,26^. As CiC promotes citrate bidirectional transport, inhibition of CiC increases intra-mitochondrial citrate levels, which could enhance TCA cycle and increase TCA intermediates, such as AKG.

The AKG is a crucial metabolite and regulator that has pleiotropic benefit, promoting heart regeneration^27^, inhibiting autophagy^28^ and ameliorating age-related osteoporosis^29^. One of AKG’s underlying mechanisms is enhancing the endogenous antioxidant pathway to alleviate oxidative stress, through conversion into glutamine, a precursor of GSH^30^. GSH, a pivotal antioxidant with multifaceted role in cellular defense mechanisms, maintains the redox state and regulate cell apoptosis.^31–33^ Our kidney metabolite analysis revealed elevated levels of AKG and GSH (Fig 5-7). Supplementation of either AKG or GSH attenuates V+G-induced oxidative stress in tubular cells and better protects against V+G-ATI in mice. Mechanistically, the protective effect of CiC inhibition requires the transport of AKG and/or glutamate across mitochondria, since knocking down either mitochondrial transporters *SLCA25A11*(AKG) or *SLC25A18* (glutamate) attenuate this antioxidant effect of CiC inhibition. Our data reinforce the critical role of oxidative stress in ATI.^34,35^ The detailed mechanisms, however, warrant further investigation.

*SLC25A11*, also known as the oxoglutarate carrier, is the primary mitochondrial transporter responsible for exchanging AKG across the inner mitochondrial membrane, thereby elevating cytosolic AKG levels.^36,37^ It facilitates the electroneutral antiport of mitochondrial AKG and cytosolic malate, a process that supports the transfer of NADH equivalents for ATP production. In addition to its metabolic role, *SLC25A11* is critical for maintaining GSH levels, thereby protecting cells from oxidative stress and cytotoxic-induced apoptosis. Recent studies have shown that loss of *SLC25A11* impairs mitochondrial GSH homeostasis, increasing vulnerability to oxidative damage and lipid peroxidation.^38,39^ Our findings demonstrate that the reno-protective effect observed in CiC KO models is mediated, at least in part, by *SLC25A11*, highlighting the pivotal role of this mitochondrial transporter in modulating ferroptosis during vancomycin-induced acute tubular injury.

Proximal tubular cells are among the most metabolically active highly enriched in mitochondria^40^, as they actively pump substrates to maintain concentration gradient and reabsorb solutes to prevent urinary loss. Proximal tubules are particularly susceptible to hypoxic stress and various forms of renal injury^41^. Under physiological conditions with adequate oxygen supply and normal mitochondrial function, proximal tubular cells primarily utilize fatty acids for oxidative phosphorylation to generate ATP. During ATI, however, there are mitochondrial injuries, reduced oxygen availability, and relative hypoxia, which decreases fatty acid oxidation. As a result, their metabolism undergoes a metabolic switch toward alternative fuels, including ketone bodies and glycolysis/glucose oxidation^42–45^. Our ^13^C flux analysis demonstrated that glucose oxidation was elevated in response to V+G-ATI. The metabolic alteration in CiC KO includes an augmented TCA cycle reserve capacity and an enhanced antioxidant.

In summary, we demonstrated that tubular-specific CiC KO induced metabolic alterations that are beneficial in enhancing endogenous antioxidant pathway (GSH, GPX4), decreasing mitochondrial ROS, ferroptosis (including lipid peroxidation and iron accumulation), hence protecting against V+G-induced kidney tubular injury. Administration of AKG or GSH recapitulated the nephroprotective effect of CiC KO. Our findings suggest the potential novel clinical application of metabolic treatment to prevent antibiotics-induced ATI.

## Data availability statement

Most raw data were generated at Johns Hopkins University School of Medicine. Stable isotope tracing data were generated at the University of Iowa, Carver College of Medicine Metabolomics Core Facility. The datasets used and/or analyzed during the current study are available from the corresponding author on reasonable request. Deposition of stable isotope tracing data to Metabolomics Workbench is pending.

## Author Contribution

**Conceptualization:** D.D. and M. H. designed the study. **Data analysis:** D.D, M.H, K.W, P.L, E.D, A.R, R.L, and H.A analyzed the data. **Writing:** M.H and D.D. wrote the paper with input from all authors. **Supervision:** E.T and D.D were involved in planning and supervision of the work.

