## Supplemental Figures for "Metabolic Reprogramming Induced by Mitochondrial Citrate Carrier Deletion Mitigates Antibiotics-Induced Acute Tubular Injury"

**Figure S1.**

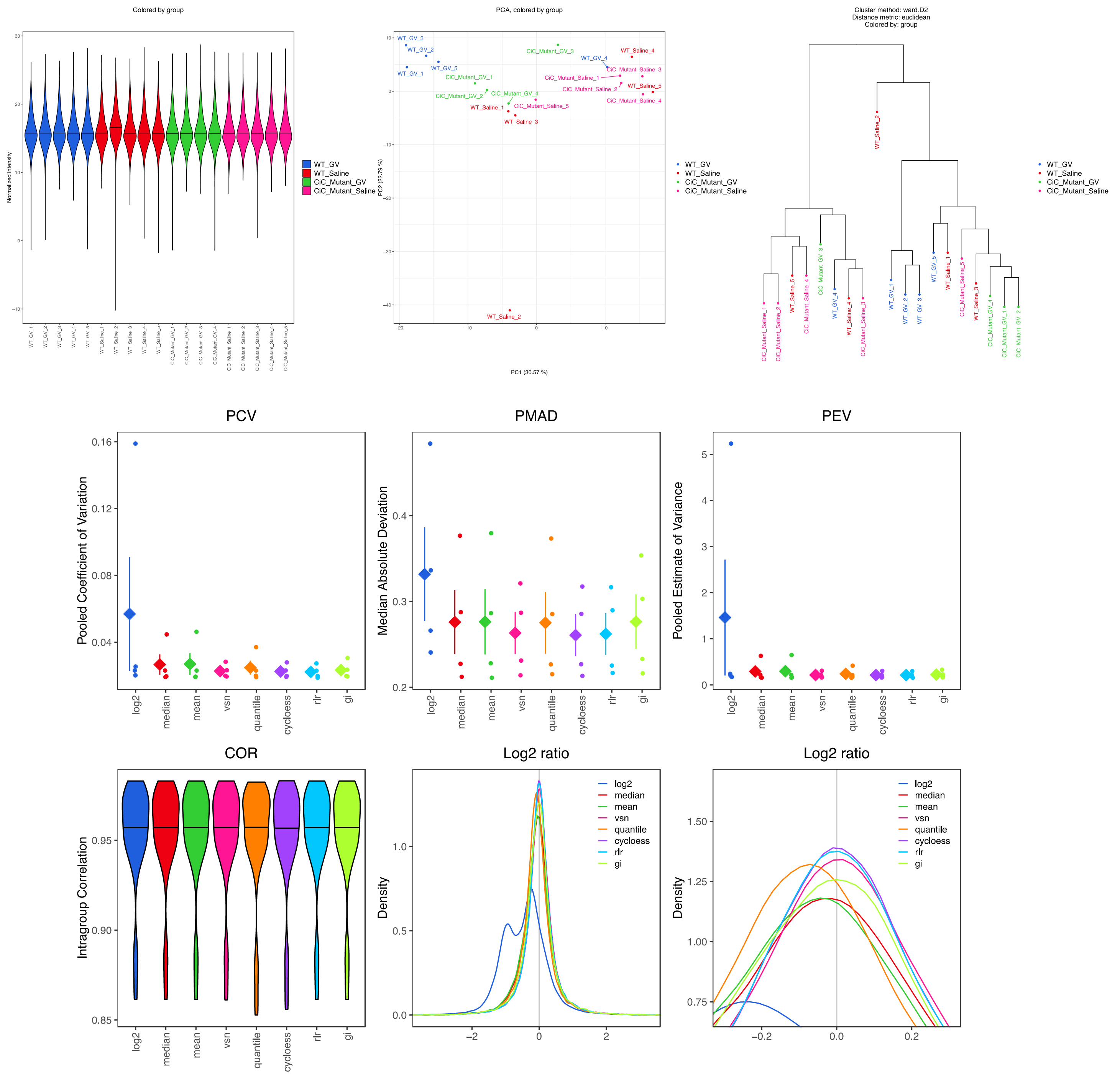
Colored by group
Normalized intensity
WT_GV
WT_Saline
CiC_Mutant_GV
CiC_Mutant_Saline
PCA, colored by group
PC1 (30.57 %)
PC2 (22.79 %)
Cluster method: ward.D2
Distance metric: euclidean
Colored by: group
PCV
Pooled Coefficient of Variation
PMAD
Median Absolute Deviation
PEV
Pooled Estimate of Variance
log2
median
mean
vsn
quantile
cycloess
rlr
gi
COR
Intragroup Correlation
Log2 ratio
Density

**Figure S1. Label-free DIA proteomics analysis of acute tubular injury in kidney tissue.**

Normalized intensity values are displayed for each biological replicate across four groups: WT saline, WT V+G, CIC-KO V+G, and CIC-KO saline. Principal component analysis (PCA) and hierarchical clustering analyses demonstrate separation and similarity among experimental groups. Quality metrics for protein quantification, including pooled coefficient of variation (PCV), median absolute deviation (PMAD), pooled estimate of variance (PEV), and intragroup correlation (COR), are plotted for each normalization method. Distribution plots of log2 ratios further assess normalization performance.

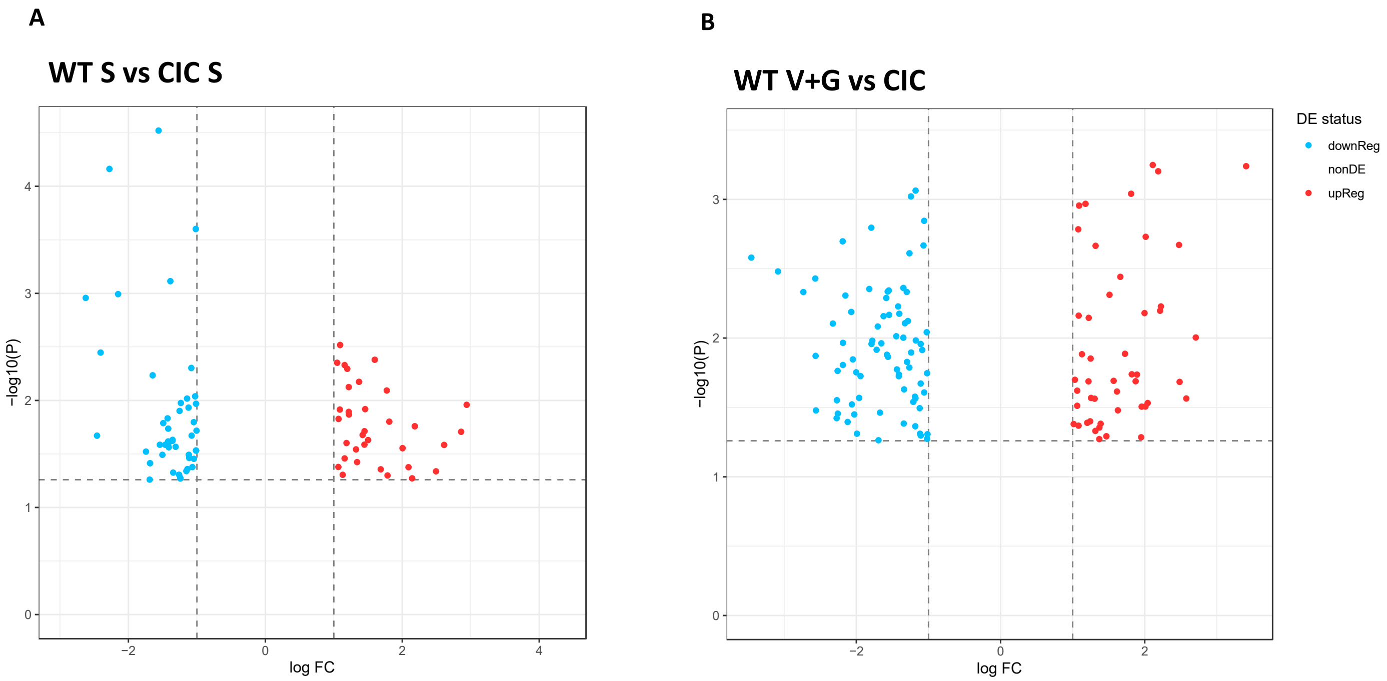


**Figure S2. Volcano plots summarize the differential protein expression analysis of the proteomics data.** (A) Wild type versus CIC-KO saline-treated control. (B) Wild type versus CIC-KO V+G-treated control (fold change > 2, $p < 0.05$).

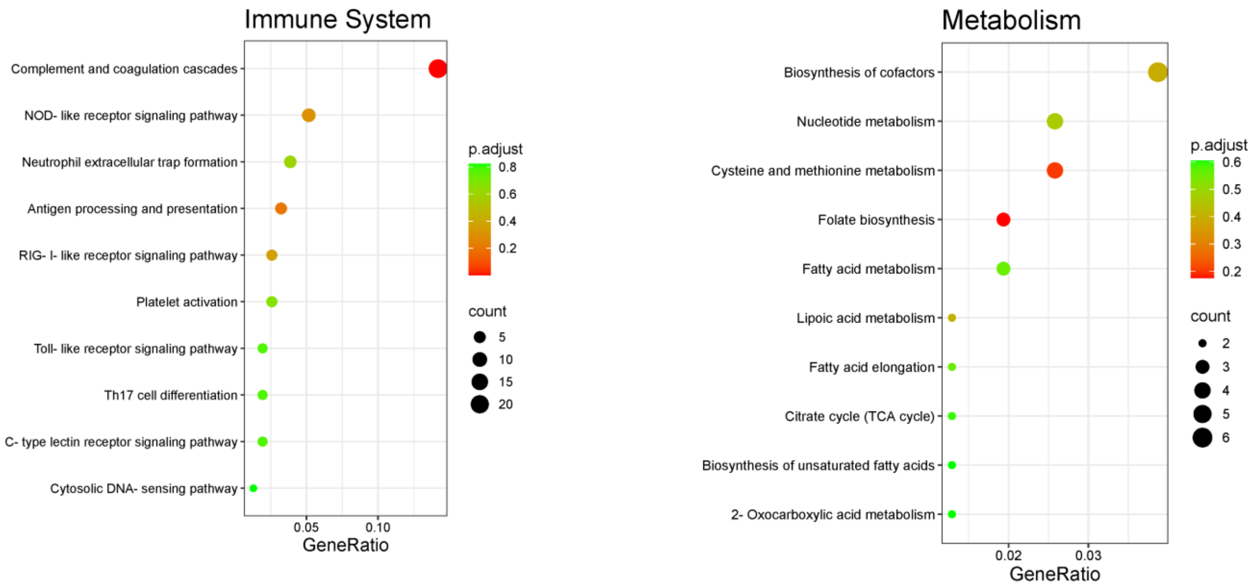


**Figure S3. KEGG pathway analysis on DIA proteomics analysis.** (A) immune system, and (B) metabolism pathway.

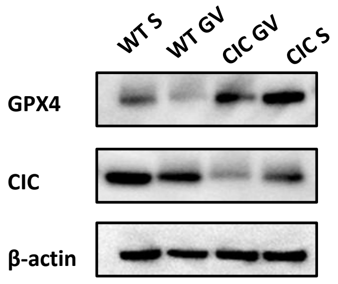


Figure S4. Representative Western blot of GPX4 in CIC KD HK-2 cells.

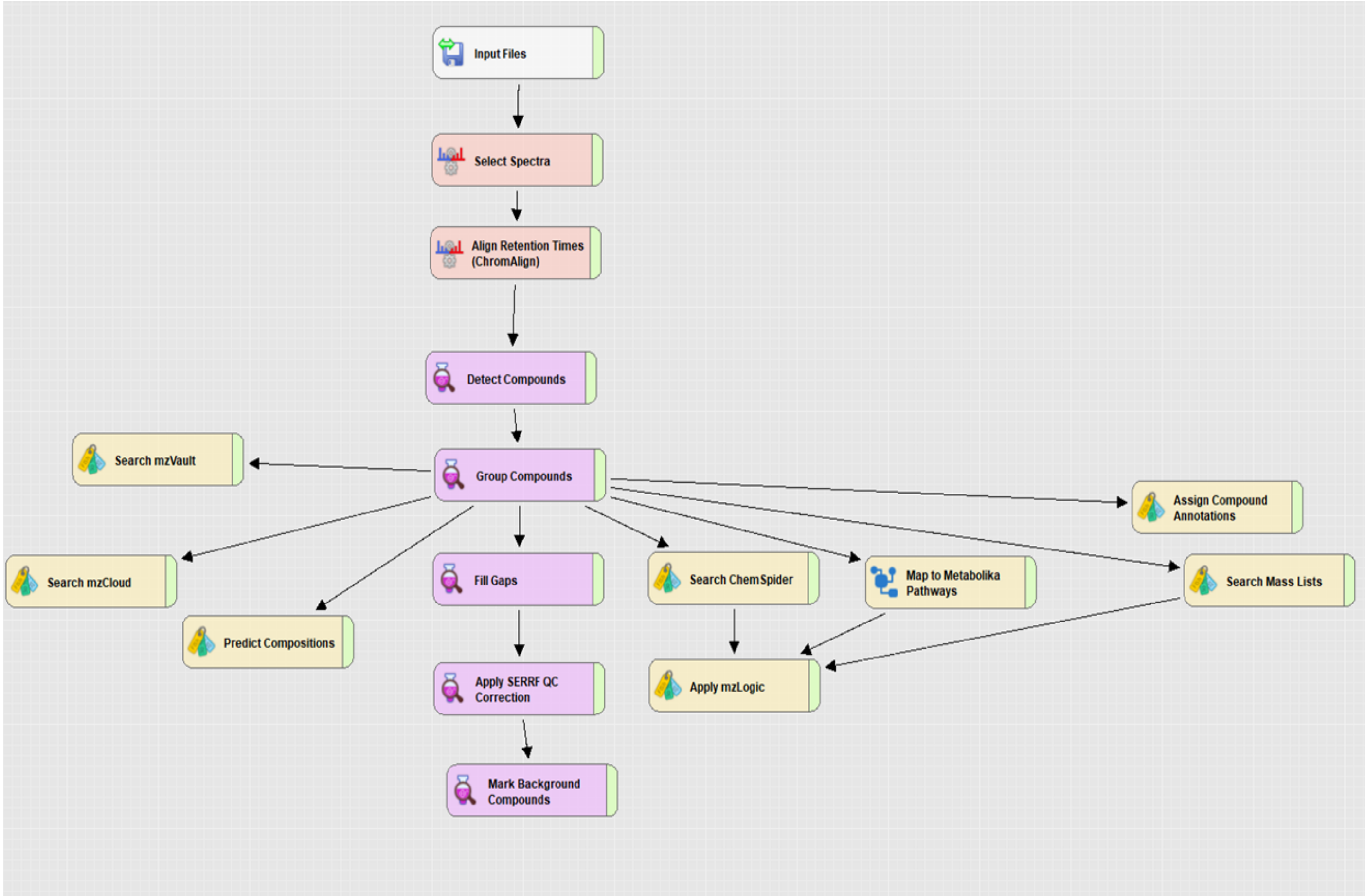


Figure S5. Workflow tree of untargeted metabolomics from Compound Discoverer 3.3.0 software, displaying data processing nodes and the associated workflow connection.

### Overview of Enriched Metabolite Sets (Top 25) Based on SMPDB

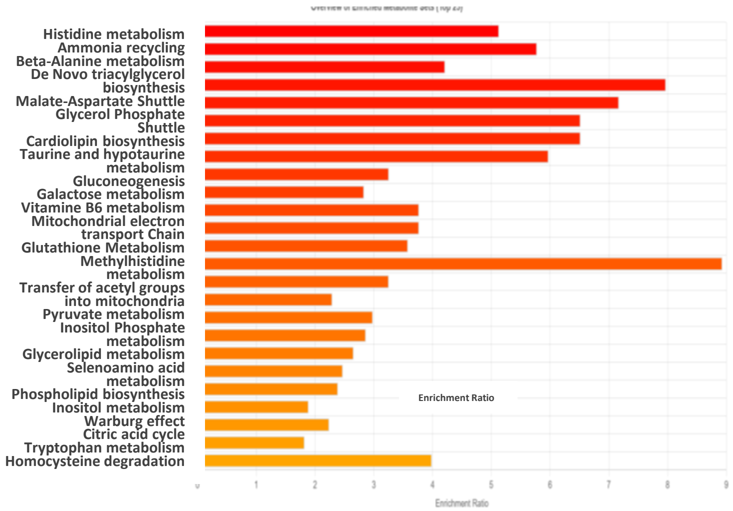


Figure S6. Metabolomics pathway analyses were performed against SMPDB.

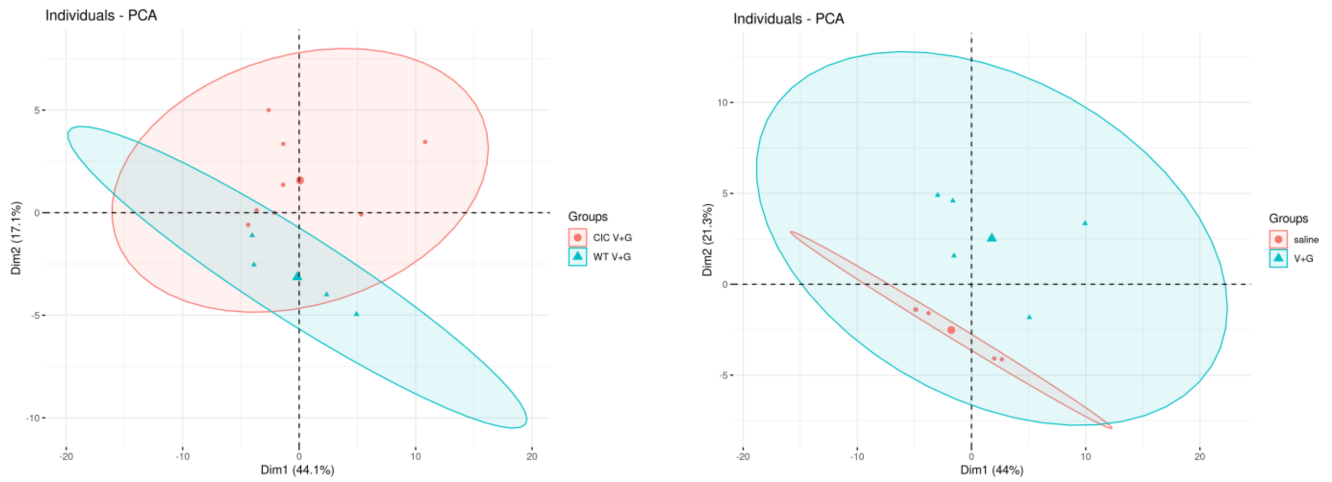


Figure S7. Principal component analysis (PCA) of metabolomics profiles highlights distinct clustering and separation between experimental groups, indicating differential proteomic signatures associated with genotype and treatment in acute kidney injury.
